# The advantages and disadvantages of short- and long-read metagenomics to infer bacterial and eukaryotic community composition

**DOI:** 10.1101/650788

**Authors:** William S. Pearman, Nikki E. Freed, Olin K. Silander

**Affiliations:** School of Natural and Computational Sciences, Massey University, Auckland, New Zealand

**Keywords:** metagenomics, Nanopore, Illumina, long read, community composition

## Abstract

**Background:** The first step in understanding ecological community diversity and dynamics is quantifying community membership. An increasingly common method for doing so is through metagenomics. Because of the rapidly increasing popularity of this approach, a large number of computational tools and pipelines are available for analysing metagenomic data. However, the majority of these tools have been designed and benchmarked using highly accurate short read data (i.e. illumina), with few studies benchmarking classification accuracy for long error-prone reads (PacBio or Oxford Nanopore). In addition, few tools have been benchmarked for non-microbial communities.

**Results:** Here we use simulated error prone Oxford Nanopore and high accuracy Illumina read sets to systematically investigate the effects of sequence length and taxon type on classification accuracy for metagenomic data from both microbial and non-microbial communities. We show that very generally, classification accuracy is far lower for non-microbial communities, even at low taxonomic resolution (e.g. family rather than genus).

**Conclusions:** We then show that for two popular taxonomic classifiers, long error-prone reads can significantly increase classification accuracy, and this is most pronounced for non-microbial communities. This work provides insight on the expected accuracy for metagenomic analyses for different taxonomic groups, and establishes the point at which read length becomes more important than error rate for assigning the correct taxon.

## Introduction

### Applying Metagenomic Methods to Quantify Community Composition

To understand ecological community diversity, it is essential to quantify taxon frequency. The most common method of quantifying taxa frequencies is through metabarcoding (Ji et al. 2013). In this method, conserved genomic regions (often 16S rRNA in the case of bacterial and archaeal species; 18S rRNA or Cytochrome c oxidase I for eukaryotic species) are amplified from the sample of interest, sequenced (most often using high-throughput methods such as Illumina), and then classified using one of several available pipelines (e.g. QIIME, MEGAN, Mothur) (Caporaso et al. 2010; Huson et al. 2016; Schloss et al. 2009). Many of these pipelines have been designed around the analysis of bacterial datasets.

In contrast to metabarcoding, metagenomic approaches do not rely on the amplification of specific genomic sequences, which can introduce bias. Instead, they aim to quantify community composition based on the recovery and sequencing of all DNA from community samples. Not only do metagenomic methods profile taxon composition in a less biased way than metabarcoding, but they can also yield insight into the functional diversity present in ecosystems (Schloss and Handelsman 2005; Keeling et al. 2014).

While metabarcoding approaches have been widely applied to both microbial and eukaryotic taxa, the vast majority of metagenomic studies have focused only on microbial communities. Unsurprisingly, the various advantages and disadvantages of using metagenomic analyses for microbial communities are well-documented (Roumpeka et al. 2017; Thomas, Gilbert, and Meyer 2012; Temperton and Giovannoni 2012). There are likely several factors driving this microbe-centric application of metagenomics, including (1) the greater level of diversity of microbial taxa; (2) the considerable number of microbial taxa that are “unculturable,” making it difficult to collect the requisite amount of DNA for genomic sequencing; (3) the availability of a multitude of non-molecular methods for quantifying multicellular taxa; and (4) the relative paucity of genomic sequence for multicellular organisms in databases (Escobar-Zepeda, Vera-Ponce de León, and Sanchez-Flores 2015) (Supp. Fig.1). This latter factor is perhaps the single largest factor in driving the bias toward microbial metagenomics.

**Figure 1.**
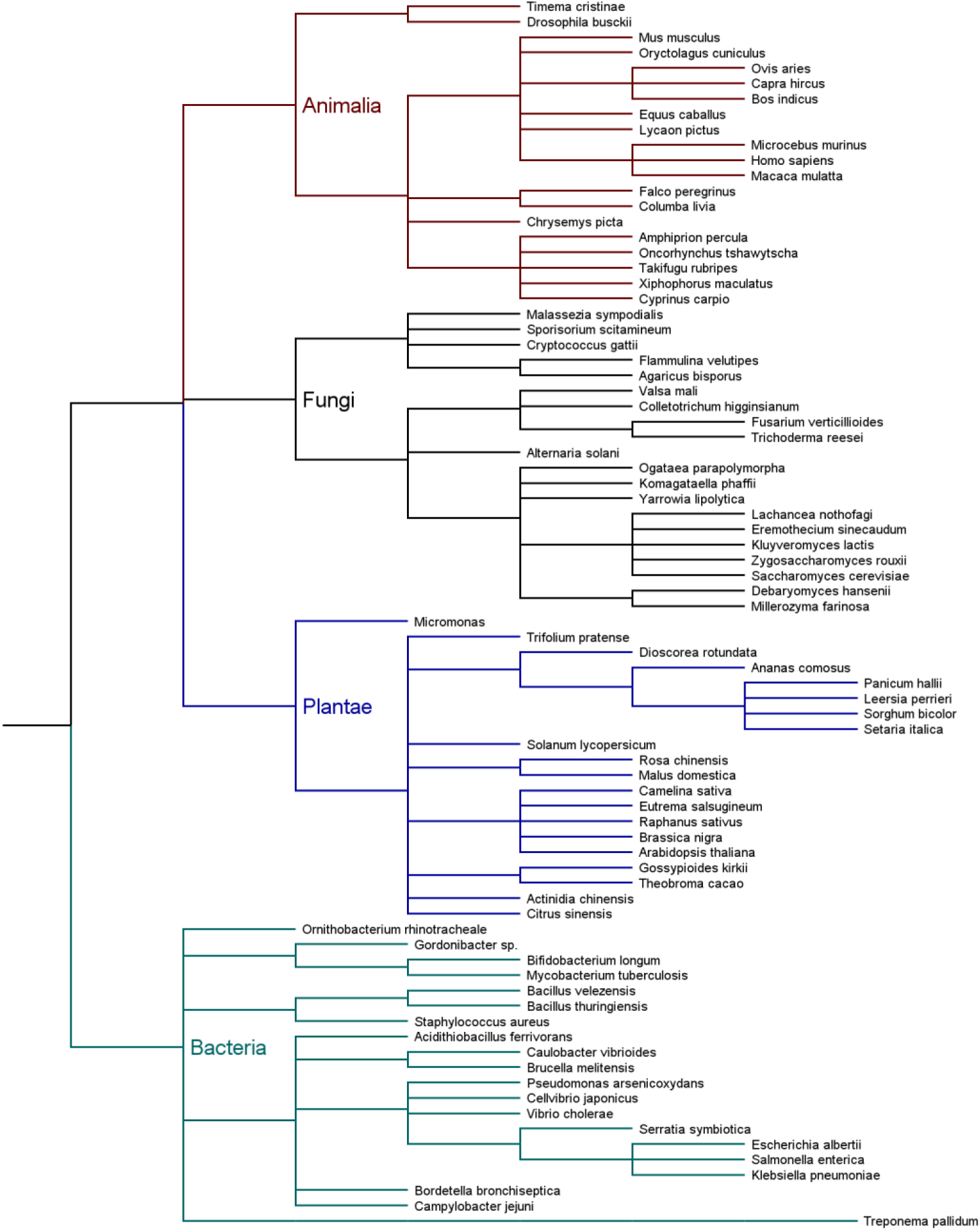
Cladogram of species included in the simulated mock community. We simulated Illumina data using dwgsim 0.1.12 (Homer 2017) with the following options: *dwgsim -e 0.0001 -E 0.0001 -N 2000 −1 100 −2 100 -r 0.0001 -R 0.01 -y 0.000 -c 0* This implements errors to mirror those in Illumina data, with constant error rates of 1e-4 and no indels (which are extremely rare in Illumina data). We generated 1,000 reads for each genome, at three read lengths: 100 bp, 150 bp, and 300 bp (a total of 240,000 reads across all taxa and lengths), and used only single end reads for all analyses.

However, the amount and diversity of eukaryotic genomic sequence data is rapidly increasing. Although multicellular metabarcode databases are currently far more complete relative to genomic databases, this gap is closing quickly. For example, the Earth BioGenome project aims to sequence the genomes of upwards of one million eukaryotic species within the next decade (Lewin et al. 2018). Regardless of the success of this effort, there are a host of ongoing eukaryotic sequencing projects, including Bat 1K (Teeling et al. 2018), Bird 10K (10,000 bird genomes (OBrien, Haussler, and Ryder 2014)), G10K (10,000 vertebrate genomes (10K Community of Scientists 2009)), and i5K (5000 arthropod genomes (Robinson et al. 2011)), among others. This suggests that within the next five years, most multicellular organisms will have at least one member of their family present in genomic databases, with some groups of multicellular organisms being completely represented at the genus level.

This would increase the utility of metagenomics for assessing membership in plant and animal communities, especially for cases in which organisms are difficult to observe or degraded. This is frequently the case for diet studies <u>(Pearman et al. 2018)</u>, many invertebrate communities such as in treeholes <u>(Gossner et al. 2016)</u> or algal holdfasts <u>(Ojeda and Santelices 1984)</u>.

### Analysis of Short-read Metagenomic Data

Many metagenomic classification analyses rely on first pass classifiers to assign reads to one or more taxa, followed by second pass classifiers that can improve on the initial classification by taking into account the number and relationship of taxa identified in the first pass. This second step often relies on a lowest common ancestor algorithm (Wood and Salzberg 2014; Kim et al. 2016; Huson et al. 2016), or by refining taxonomic representation by examining the results from the first pass classifier (Lu et al. 2016).

The most widely used first pass classifier is BLAST, and it is considered gold standard (McIntyre et al. 2017). However, BLAST is not computationally efficient enough to deal with tens or hundreds of millions of reads. Thus, algorithms for fast metagenomic classification have been the subject of intense research over the last few years, and include k-mer based approaches such as CLARK (Ounit et al. 2015), Kraken and related tools (Kraken, Kraken2, and KrakenUniq) (Wood and Salzberg 2014), Centrifuge (Kim et al. 2016), EnSVMB (Jiang et al. 2017), and Kaiju (Menzel, Ng, and Krogh 2016), as well as reduced alphabet amino acid based approaches such as DIAMOND (Buchfink, Xie, and Huson 2015). In almost all cases these have been designed and benchmarked using short read data (McIntyre et al. 2017).

### Analysis of Long-Read Data Metagenomic data

The advent of “third generation” single molecule long read technologies (PacBio and Oxford Nanopore) has significant implications for metagenomic analyses, most notably for genome assembly (Frank et al. 2016; Nicholls et al. 2019). These technologies allow read lengths of 10 kilobase pairs (Kbp) and beyond, in strong contrast with the approximately 300 base pairs (bp) limit of Illumina. However, both PacBio and Nanopore technologies have far higher error rates (88-94% accuracy for Nanopore (Wick, Judd, and Holt 2018) and 85-87% for PacBio (Ardui et al. 2018)). The lower accuracy of Nanopore and PacBio (non-circular consensus) sequence reads may affect the success of current classification methods, and there are few algorithms designed to exploit long-read data.

As a first approach toward determining the use of long-read technologies for metagenomic applications, we would like to understand the relative advantages and disadvantages of using short accurate reads versus long error-prone reads. Recent work has shown that relatively high genus level classifications of approximately 93% have been achieved using Nanopore-based metagenomic analyses of a mock bacterial community (Brown et al. 2017). Here we expand this analysis to allow direct comparison between short and long read approaches. In addition, we compare metagenomic classification success in microbial communities as compared to communities of multicellular organisms. We find that longer reads, despite their higher error rate, can considerably improve classification accuracy compared to shorter reads, and that this is especially true for specific taxa.

## Methods

### Genomic data

For each of four major taxonomic divisions (bacteria, fungi, animals, and plants), we downloaded 20 genomes from GenBank (Benson et al. 2013). Within each of these divisions, we included genomes from a total of 22 classes, 46 orders, and 58 families (Figure 1).

### Read simulation

We simulated Nanopore reads using NanoSim 2.0.0 (Yang et al. 2017) with the default error parameters for *E. coli* R9 1D data. This method uses a mixture model to produce simulated reads with indel and error rates similar to real datasets. The error model is applied equally to all parts of a read, and the read lengths are drawn from a distribution approximating real data. To create simulated read data of specific lengths, we truncated the simulated reads after the relevant number of basepairs using a custom perl script (i.e. to simulate 100bp Nanopore reads, we truncated all reads in a simulated dataset to 100bp). We did this for read lengths varying from 100 bp to 4,000 bp at 100 bp intervals, simulating 1,000 reads per interval for all taxa (a total of 40,000 reads for each taxon, and 3.2 million reads for all taxa and read lengths).

### Sequence Classification

We used BLAST 2.7.1 (Madden 2013) and Kraken2 (Wood and Salzberg 2014) for sequence classification. We created a local custom database consisting of the NCBI nt database (downloaded on Feb 8 2019) and the genomes of the 80 taxa that we used to test classification success. We used the default alignment parameters for BLAST, except for implementing a maximum e-value of 0.1. We used the match with the highest bit score for all downstream analyses. For Kraken2 analyses we used the default parameters (in which the k-mer length is 35 bp and default minimiser length is 31 bp). For Kraken2 we used the taxon assigned by the lowest common ancestor (LCA) algorithm employed in Kraken2.

### Accuracy metrics

To assess the effects of read length on classification accuracy we focus our analysis only on how often a read is assigned to the correct taxon. For our simulated reads there are three possible outcomes when querying a database (Table 1).

**Table 1.** Description of outcomes for database queries.

| Description of outcome | Metric | Notation |
| --- | --- | --- |
| A read query from taxon A returns a match from taxon A | True match (we correctly infer taxon A is present) | $M_{\text{true}}$ |
| A read query from taxon A returns a match from a taxon that is not A | False match (we infer taxon A is absent due to a secondary match) | $M_{\text{false}}$ |
| A read query from taxon A returns no hit at all | Failed query (we infer taxon A is absent due to database paucity). | $M_{\text{fail}}$ |

We expect that taxa that are well represented in the database, and which have few closely related taxa, will have high rates of true matches. Taxa with many close relatives in the database will have many false matches. Taxa that are poorly represented in the database will have high rates of failed queries. Both of these latter results are in a class usually referred to as false negatives: we falsely infer taxon A is absent. However, they largely arise from different mechanisms. Importantly, as genomic databases become more complete, we expect the fraction of failed queries will decrease. At the same time we expect that the fraction of false matches may increase, as more and more closely related taxa become present in the database. The exact nature of this tradeoff is not well explored. Novel statistical approaches, such as Bayesian re-estimation of species frequencies, may mitigate the problem (Lu et al. 2016); however, improved methods are required address this problem (Nasko et al. 2018).

There are other aspects of classification success that we do not focus on here. The first of these is the notion of a true negative: a sequence that is known to *not* arise from any taxa, should not return a match to any taxa. This is not a biologically realistic situation (all sequences arise from a taxon), although this aspect is useful when trying to assess the performance of different classifiers [ref Gardner] and presenting the full truth table. The second aspect we do not consider here are false positives: if a read query matches taxon A, but does not arise from taxon A. We would thus falsely interpret taxon A as being present in a community. This metric is intrinsic to the composition of the community rather than just each taxon and the database. For example, if taxon A dominates the community, then it cannot have high rates of false positives relative to true positives simply because the vast majority of read queries from the community will be from taxon A and thus true positives. Conversely if taxon B is extremely rare, there will be a large number of false positives relative to true positives, as very few read queries will be from taxon B, resulting in a very small number of true positives.

Thus, we use a simplified set of metrics (see Table 1) that are not intrinsically related to community composition: true matches, false matches, and failed queries. We used our simulated genomic sequence reads from 80 taxa to quantify these three outcomes at both the genus and family level. To assign genus and family from species, we used the NCBI taxonomy database (Federhen 2012) (which is used by BLAST as the default taxon classifier).

We calculate two ratios from the three metrics in Table 1. The first is the fraction of true positives classified correctly (i.e. recall):

Recall = M_true_/(M_true_ + M_false_ + M_fail_)

The second is the ratio of true matches to false matches. This simply excludes failed queries from the equation. We term this second metric classification success.

Classification Success = M_true_/(M_true_ + M_false_)

The critical difference between these metrics is that taxa which are poorly represented in the database may nevertheless have high rates of classification success, although recall will necessarily be low. However, as the fraction of failed queries approaches zero (which we expect as genomic databases grow), these two metrics become equivalent.

## Results

We first looked only at short read lengths to quantify the effects of sequencing technology and classifier (BLAST or Kraken2) on recall at the level of genus. For both bacteria and fungi, we found that recall was at or above 99.9% for Illumina reads of any length (100bp, 150bp, or 300bp), for both BLAST and Kraken2 (Fig. 2). In strong contrast, for Nanopore data, recall was far lower; approximately 25% for 100bp reads and increasing to 75% at 300bp. In general, Kraken2 had slightly lower recall than BLAST.

**Figure 2.**
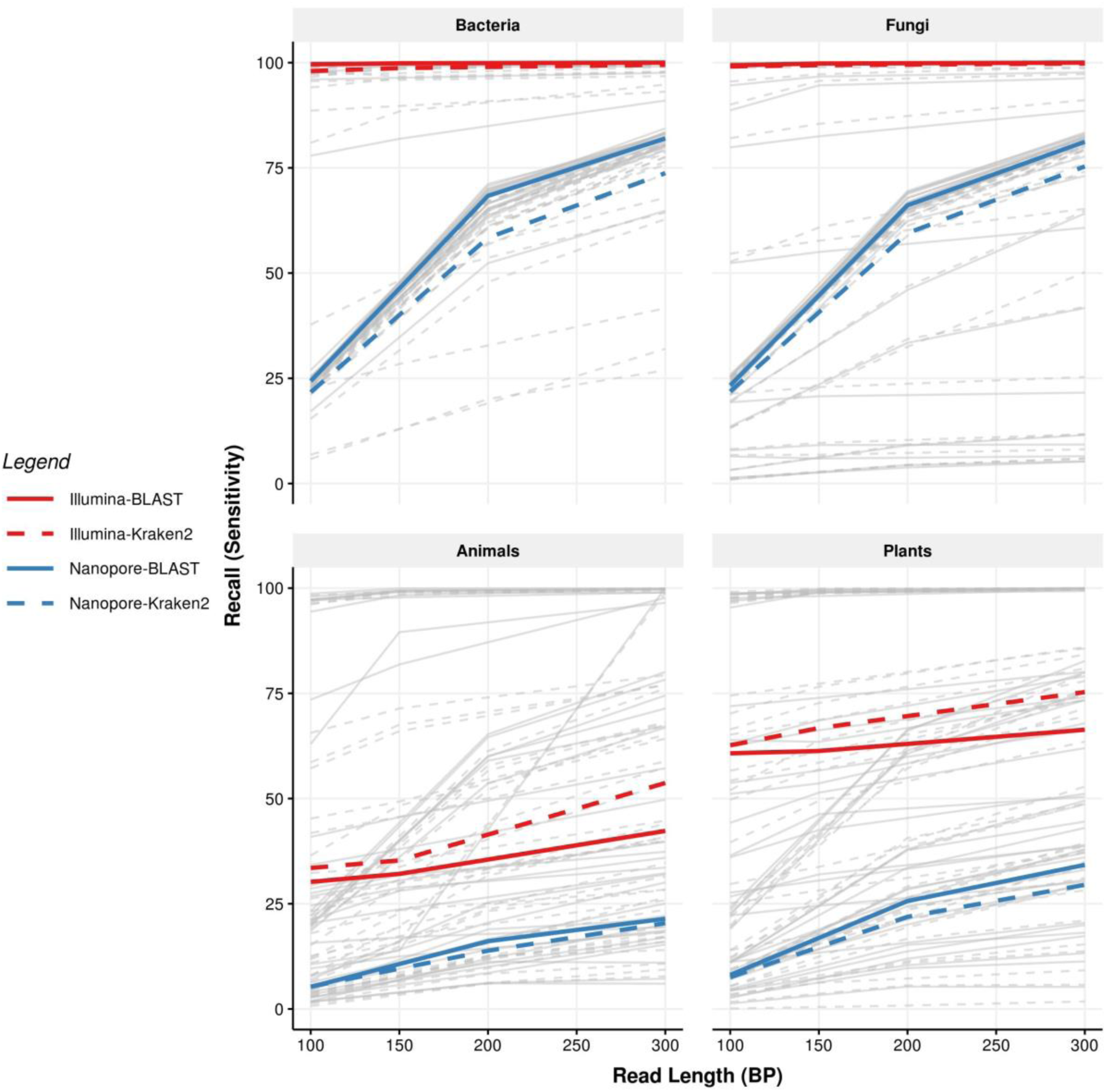
Recall is consistently higher in bacteria and fungi than plants or animals for both short Illumina and Nanopore reads. Each panel shows recall for the different kingdoms. Recall for individual taxa is indicated in grey, with median recall shown by dashed (Nanopore) or solid (Illumina). Blue lines indicate recall rates for reads classified using Kraken2; red for reads classified using BLAST. Illumina reads exhibit consistently higher recall; bacteria and fungi exhibit higher recall than plants or animals.

However, for plants and animals, average recall was low regardless of sequencing technology. Average recall for Illumina reads peaked at approximately 55% and 75% for animals and plants, respectively (Fig 2, light blue lines). Nanopore recall rates peaked at just over 20% and 35% for animals and plants, respectively. However, this was highly taxon-dependent, with some taxa consistently having recall near 100%, while others remained close to 0% regardless of sequencing technology or read length (Fig. 2, grey lines). Perhaps surprisingly, on average Kraken2 outperformed BLAST for Illumina reads for both plant and animal taxa.

We next quantified differences in classification success (the proportion of all classified reads that were correctly classified), again considering only short read lengths. For bacteria and fungi, both Illumina and Nanopore reads exhibited high classification success, with the exception of Kraken2 classification of Nanopore reads (Fig. 3). For each sequencing method and classifier, classification success for plants and animals was low relative to bacteria and fungi. For both Illumina and Nanopore, BLAST resulted in approximately 87% and 97% of reads being correctly classified, for animals and plants respectively. However, Kraken2 success was far lower, especially for Nanopore reads, peaking at 54% in animals (Fig. 3). Over this range of read lengths, we found only a weak relationship between read length and classification success, in contrast to the results for recall.

**Figure 3.**
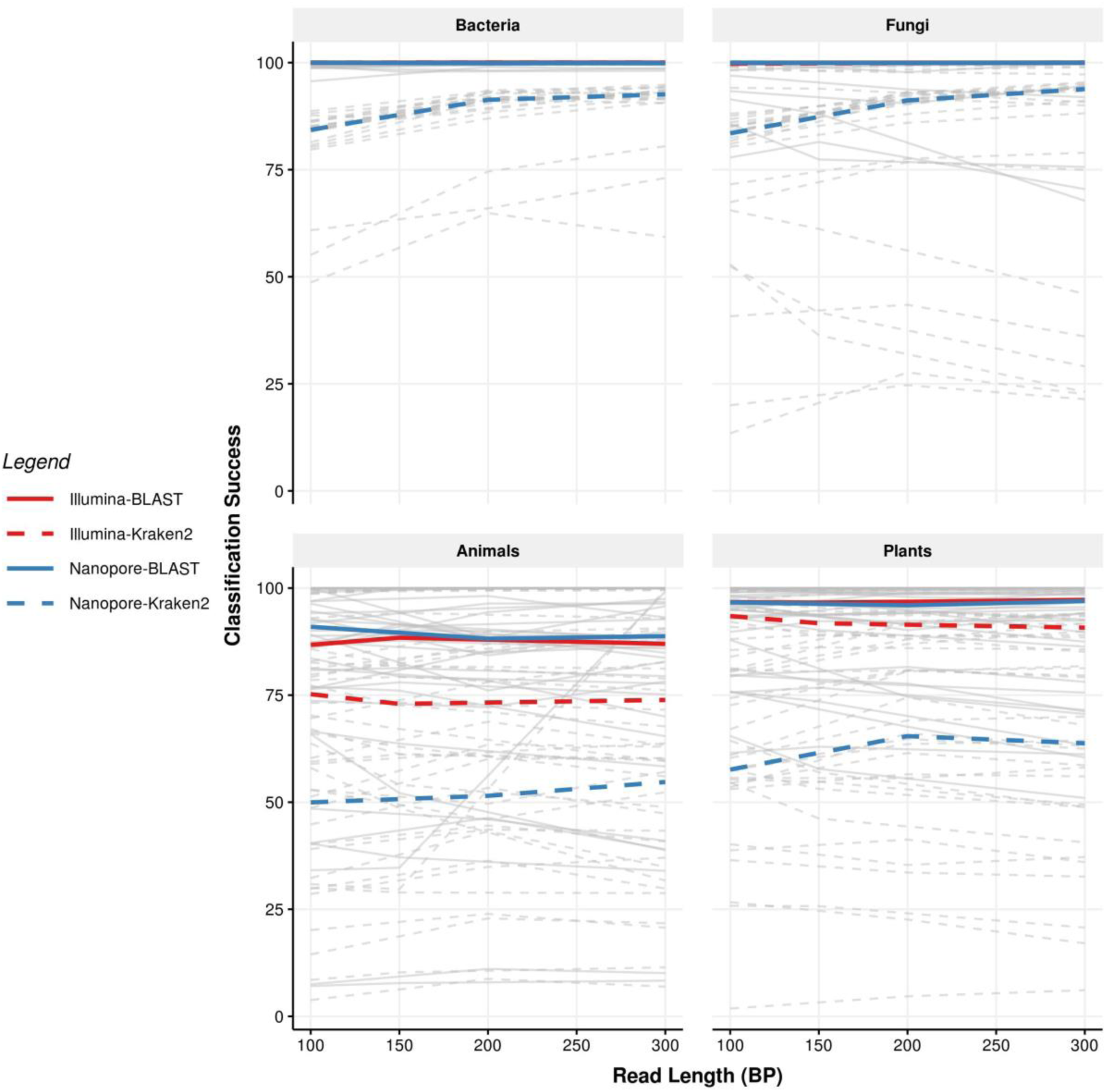
Classification success for short reads is weakly related to read length and strongly dependent on classification method. Each panel shows recall for the different kingdoms. Classification success for individual taxa is indicated in grey, with median classification success shown by solid lines (Illumina) or dashed lines (Nanopore). Blue lines indicate recall rates for reads classified using Kraken2; red for reads classified using BLAST. For bacteria and fungi, median classification rates of Illumina-BLAST, Illumina-Kraken2, and Nanopore-BLAST are almost exactly 100% for all read lengths.

It is perhaps expected that highly accurate Illumina reads would result in more accurate taxonomic classification than long error-prone Nanopore reads. However, it is possible to obtain Nanopore reads far in excess of 300bp (reads up to 2 megabase pairs have been sequenced), so we next quantified recall and classification success for reads with lengths up to 4,000 bp. Because such read lengths are not currently possible to obtain using Illumina technology, we did not measure recall and classification success for Illumina reads of similar lengths.

We observed similar relationship between read length and recall for both BLAST and Kraken2. For bacteria and fungi, recall increased from ∼20% using 100 bp reads to almost 100% when using 1500 bp reads. For animals and plants we observed similar trends, although at no point did recall approach 100%. However, long Nanopore reads surpassed the recall of even the longest Illumina reads (300 bp) classified with Kraken, with crossover points at approximately 3000 bp for animals and 2500 bp for plants (Fig. 4, red and blue solid lines).

**Figure 4.**
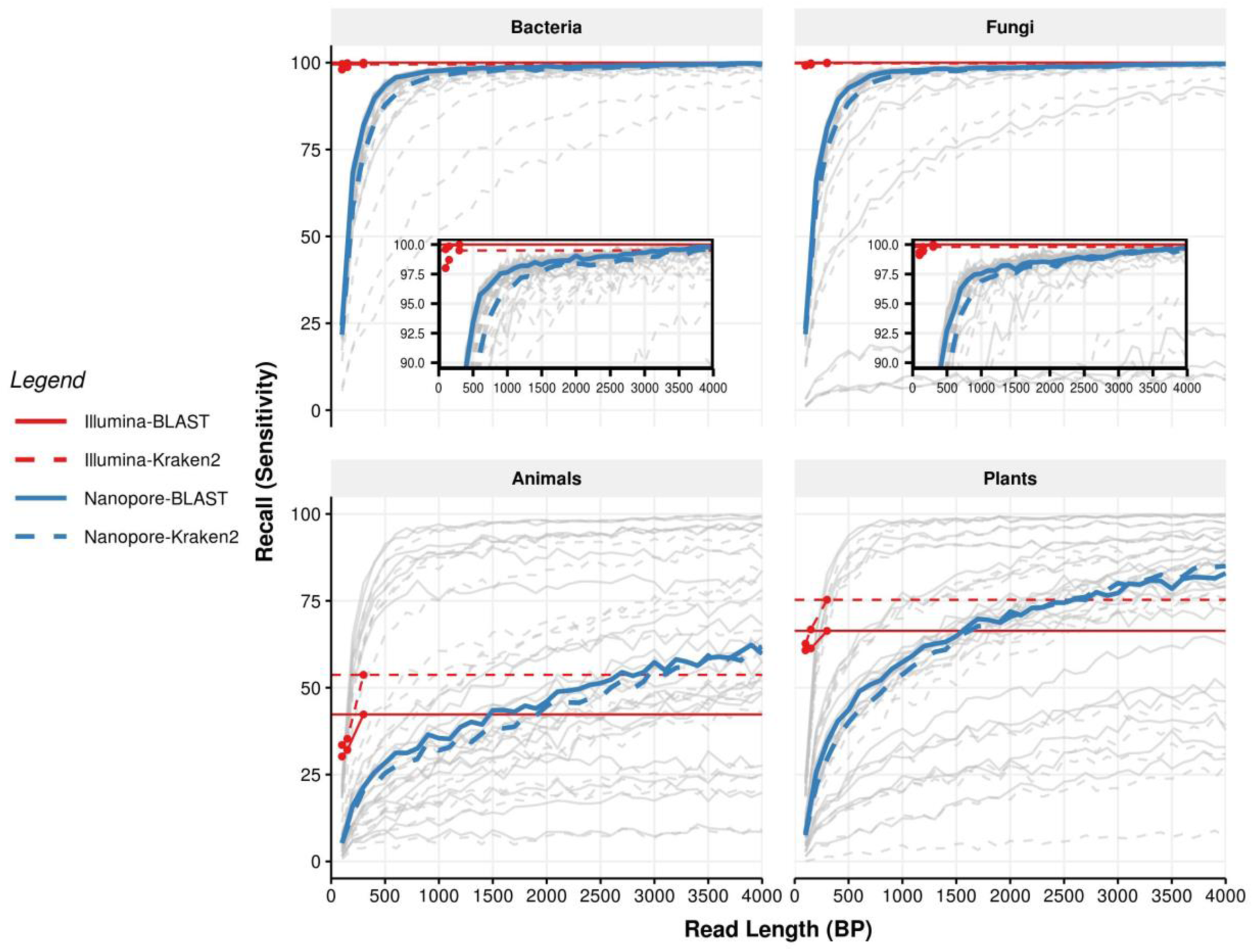
Long Nanopore reads equal or surpass the recall of the longest Illumina reads for both BLAST and Kraken2. Each panel shows recall for the different kingdoms. Recall for Nanopore reads for individual taxa is indicated in grey, with median recall indicated by dashed lines, either blue (Kraken2) or red (BLAST). The recall rates for 300 bp Illumina reads are shown as thin solid lines, again either blue (Kraken2) or red (BLAST). Coloured points show the recall for all Illumina reads of all lengths (100 bp, 150 bp, and 300 bp).

We also considered this metric at the level of family. In this case found that for animals, Nanopore reads surpassed Illumina reads only at lengths close to 4000 bp, reaching approximately 70% recall at this point (Supp Fig. 2). However, for plants Nanopore recall surpassed Illumina recall at 2500 bp, with 4000 bp reads yielding a recall of approximately 90%. We found again that for both animals and plants, Kraken2 recall surpassed BLAST when relying on Illumina reads.

We next examined classification success at longer read lengths. For BLAST we observed no relationship between classification success and read length for any taxon (Fig 5.). Bacteria and fungi both had consistently high classification success (median 100%), while animals and plants had lower classification success (median 82% and 96%, respectively). However, for Kraken2 we observed a consistent increase in classification success as read length increased. However, this never exceeded the classification success we observed for BLAST, nor did it succeed the classification success we observed for short accurate Illumina reads.

**Figure 5.**
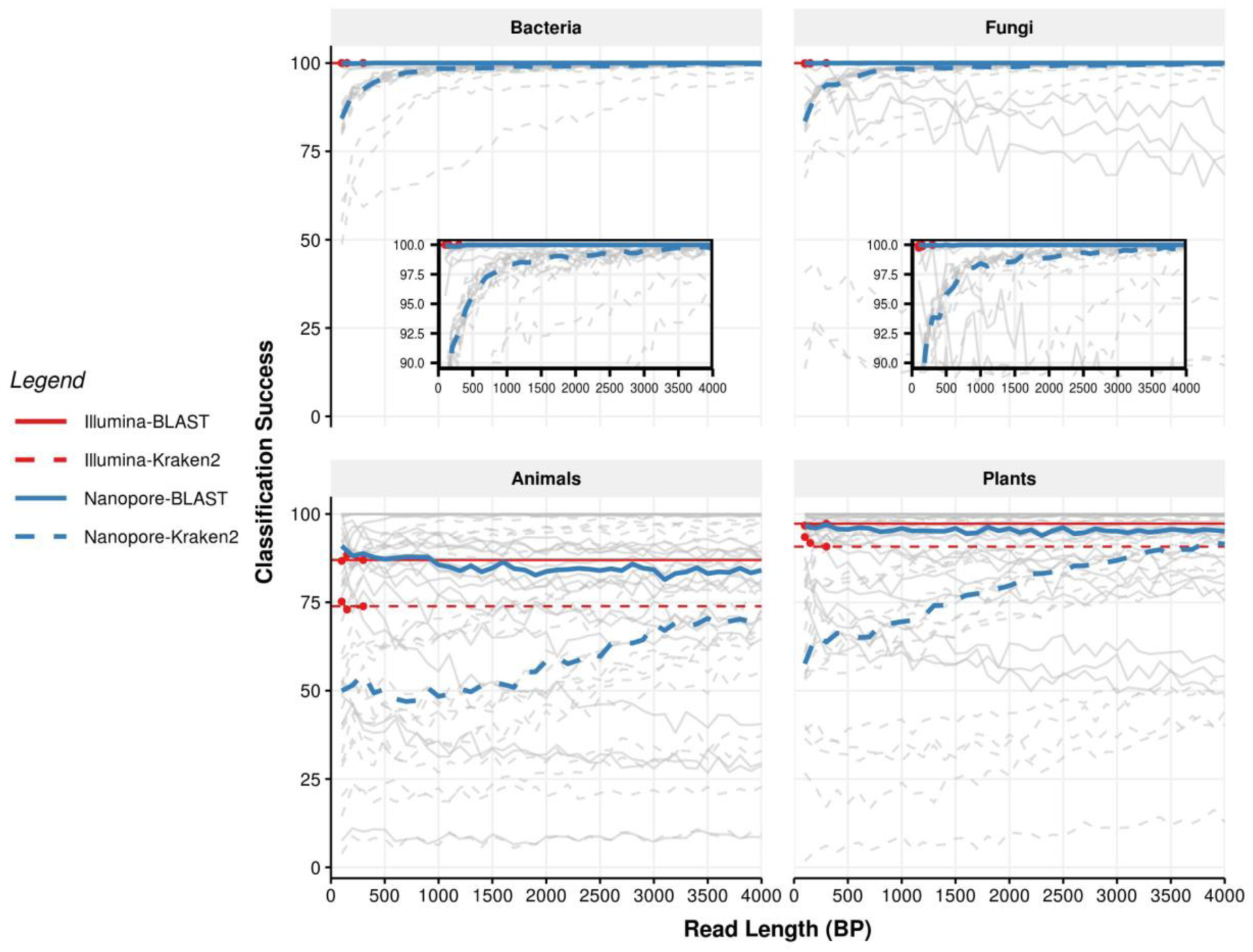
Classification success for long reads is dependent on read length only for Kraken2 classification. Each panel shows classification success for the different kingdoms. Classification success for individual taxa is indicated in grey, with median classification success shown by solid lines (Illumina) or dashed lines (Nanopore). Blue indicates classification success rates for reads classified using Kraken2, while red indicates those classified using BLAST. For animal and plants, the classification success of Kraken2 depends strongly on read length, and never surpasses BLAST or Illumina at any length.

Finally, we tested classification success at the level of Family. In this case, we observed that for BLAST, the classification success for plants was approximately 99% overall read lengths, while for Kraken2 only 4000 bp reads reached this level. For animals, BLAST classification success was approximately 95% over all read lengths, but for Kraken2 reached a maximum of 85% at the longest read lengths (Supp. Fig. 3).

## Discussion

Here we have compared the relative accuracy of taxon classification using simulated short accurate reads (Illumina) and long, error-prone reads (Nanopore) with known ground truth. We have used two simple metrics of success: recall (the ratio of correctly classified reads to all reads) and classification success (the ratio of correctly classified reads to all classified reads). We have tested taxon classification using a broad range of taxa, including bacteria, fungi, animals, and plants.

Recall for both BLAST and Kraken2 was improved by the use of long reads, especially in the case of animals and plants, for which recall improved almost three-fold as read length increased from 300 bp to 4,000 bp. Generally both Kraken2 and BLAST achieved similar levels of recall. The exception was for short reads for animals and plants, for which Kraken2 was more accurate than BLAST.

We found no relationship between classification success and read length for BLAST. This implies that the ratio of correctly classified reads to all classified reads remains relatively constant over different read lengths. However, the number of reads that are classified *at all* increases with read length (causing an increase in recall). These observations are in line with what has been observed by others (McHardy et al. 2007). The exception to this lack of relationship between classification success and read length was for Kraken2, for which the proportion of correctly classified reads increases with read length by more than 50% for both plants and animals.

Our results also indicate that recall for long Nanopore reads was equal to or higher than short Illumina reads. This was true regardless of kingdom, or classification method, with Nanopore surpassing 300 bp Illumina reads at approximately 1500 bp for plants and animals, and surpassing 150 bp Illumina reads at between 1500 bp and 3000 bp for bacteria and fungi, depending on the methodology (Fig. 4). Even the longest Illumina reads, at 300 bp, were outclassed by Nanopore at between 3500 and 4000 bp, depending on methodology. These results do suggest that one approach to improve Nanopore classification accuracy is to impose minimum read lengths. This can be achieved by performing size selection during library preparation or during computational analyses.

At first glance, then, there appears to be a clear trade-off between short read Illumina and long read Nanopore sequencing for metagenomic analyses. While Nanopore allows higher recall at long read lengths, this advantage is offset by the fact that Illlumina generally provides more reads per run. At most, recall for Nanopore improves 50% beyond 300 bp Illumina reads, while classification success is similar (using BLAST). Thus, if the read capacity of Illumina runs is 50% or more than Nanopore, the number of classified reads will be maximised using Illumina technology - on a per sequencing run basis. However, for many researchers the more relevant metric is cost per read. In this case, MinION read yields are approximately equal to MiSeq, and only HiSeq or NovaSeq provides a clear cost advantage over Nanopore MinION. On the other hand, cost per read for PromethION are not far from NovaSeq. Thus, we find no clear advantage in using Illumina over Nanopore given the observed classification accuracy for long inaccurate Nanopore reads.

### Differences in accuracy between bacteria, fungi, animals, and plants

We find very large differences in classification accuracy (mostly in terms of recall) for bacteria and fungi versus plants and animals. The discrepancy between taxonomic groups likely arises from a variety of factors. Among these are the higher degree of divergence between bacterial species relative to animal and plant species, and the complexity of bacterial genomes compared to eukaryotic genomes. We discuss these factors below.

Bacterial taxa are often considered separate species once they have diverged by 6% ANI (Average Nucleotide Identity) on a genomic level (Stackebrandt and Goebel 1994; Konstantinidis and Tiedje 2005). The degree of nucleotide divergence between eukaryotic species is not standardised (Cognato 2006), and species are generally designated as such based on the biological species concept put forward by Mayr (Mayr 1999). Although divergence levels differ substantially between loci (as for bacteria), for some loci general ranges for eukaryotic species have emerged. For example, for mitochondrial COI, between-species divergence is usually greater than 3% (Song et al. 2008; Lefébure et al. 2006). These loci are among the fastest diverging loci in plant and animal genomes, and many other loci may differ by far less than 1% between species. Due to this low level of divergence, metagenomic classifiers may frequently classify animal and plant genera with lower accuracy than bacterial genera.

A second explanation for the increased classification success in bacteria and fungi is that these genomes contain fewer repetitive elements than animals or plants (Treangen et al. 2009). Although such repetitive regions are usually masked from classifiers (including BLAST and Kraken2), this masking may not be complete.

A third reason is that the genomic databases for plants and animals are far less complete than for bacteria and fungi. There is a large difference in the number of genomes and sequences available for different Kingdoms, with bacteria having significantly more species present than the next closest kingdom (See Supp. Fig.1). However, we expect this factor will be mitigated in the future as genomic databases continue to expand and computational search methods continue to improve.

### Differences in accuracy between Kraken2 and BLAST

We observed similar levels of recall for BLAST and Kraken2 over most reads lengths. However, there were strong differences in classification success. For short reads, Kraken2 classification success was far lower than BLAST. As read lengths increased, Kraken2 classification success approached BLAST. Part of this is likely due to longer reads allowing multiple k-mer matches, decreasing the probability of a false positive classification. One perhaps underappreciated advantage of Kraken2 over BLAST is that Kraken2 has reduced sensitivity to structural variation within reads. As Kraken2 allows multiple k-mers to match within a read, structural changes (e.g. inversions) are less likely to influence the outcome of Kraken2 matching. Such structural changes may influence BLAST due to the matching and extend algorithm. Thus for long reads, classifiers that are insensitive to synteny may be more successful, especially for taxa in which structural rearrangements are common.

### Conclusions

Here we have shown despite being error-prone, Nanopore reads are useful for metagenomic classification due to their increased length, and that for plant and animal communities, the classification accuracy of long Nanopore reads exceeds that of Illumina. We found that classification accuracy is more dependent on the set of taxa being considered than on the metagenomic classifier being used (Kraken2 or BLAST), and that this was true for both short accurate (Illumina) and long error-prone (Nanopore) sequence data. Together these data suggest that one consideration in selecting a metagenomic sequencing method (i.e. long or short read) is the taxonomic group of interest.

## Declarations

### Ethics approval and consent to participate

not applicable

### Consent for publication

approved by all authors

### Availability of data and material

data for this manuscript will be uploaded to DataDryad if manuscript is accepted

### Authors’ contributions

WP, NF, and OS conceived the project, WP and OS simulated and generated the data. WP and OS analysed the data. WP, NF, and OS wrote the paper.

## Acknowledgements

Thanks to Paul Gardner for his helpful and insightful comments on the manuscript.

## Supplementary Materials

**Supplementary Figure 1.**
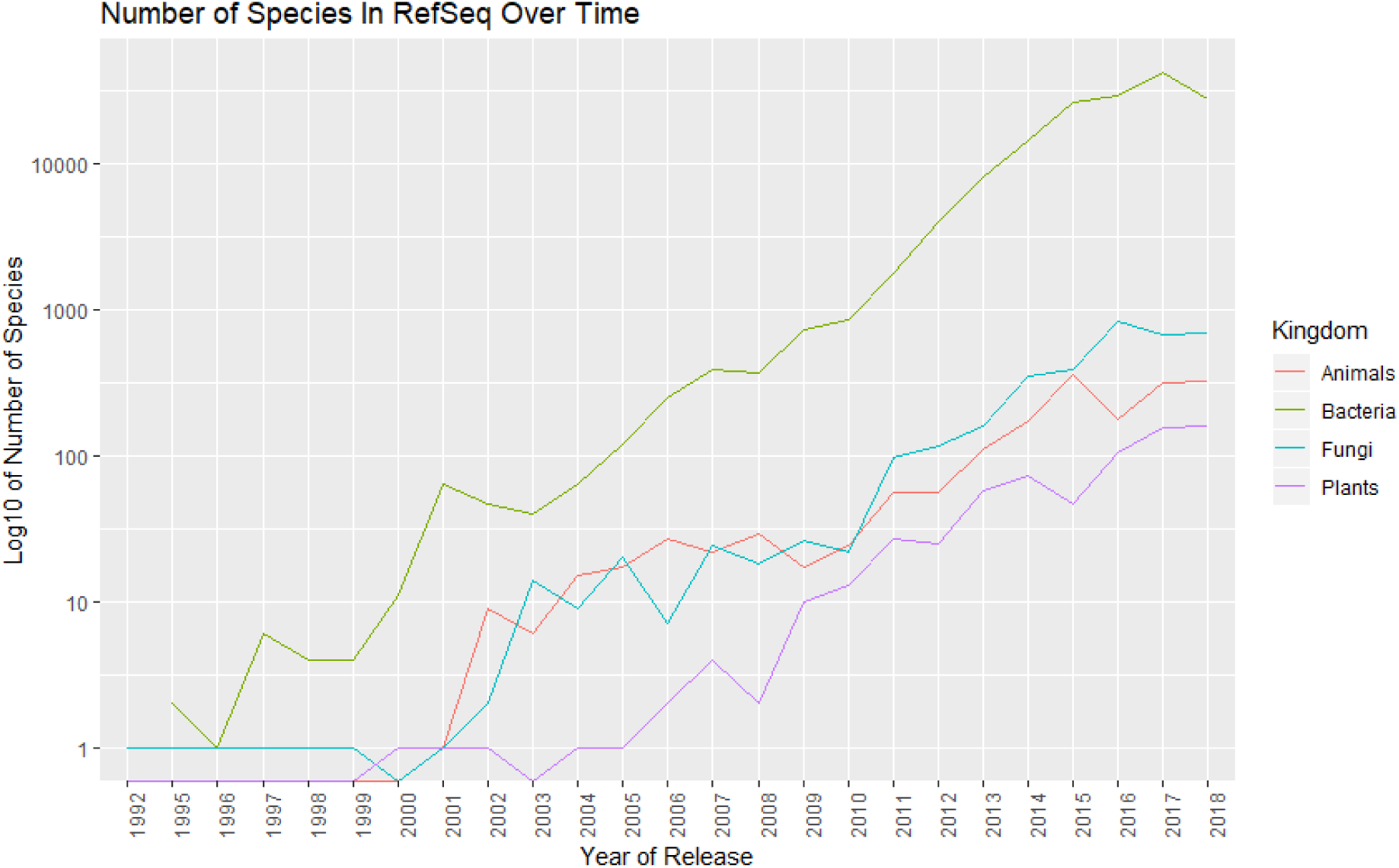
The number of species present in the NCBI RefSeq database has grown roughly exponentially over time. Note that the y-axis is plotted on a log scale. Data were retrieved from the RefSeq database (O’Leary et al. 2016): https://ftp.ncbi.nlm.nih.gov/genomes/GENOME_REPORTS/eukaryotes.txt and https://ftp.ncbi.nlm.nih.gov/genomes/GENOME_REPORTS/prokaryotes.txt

**Supplementary Figure 2.**
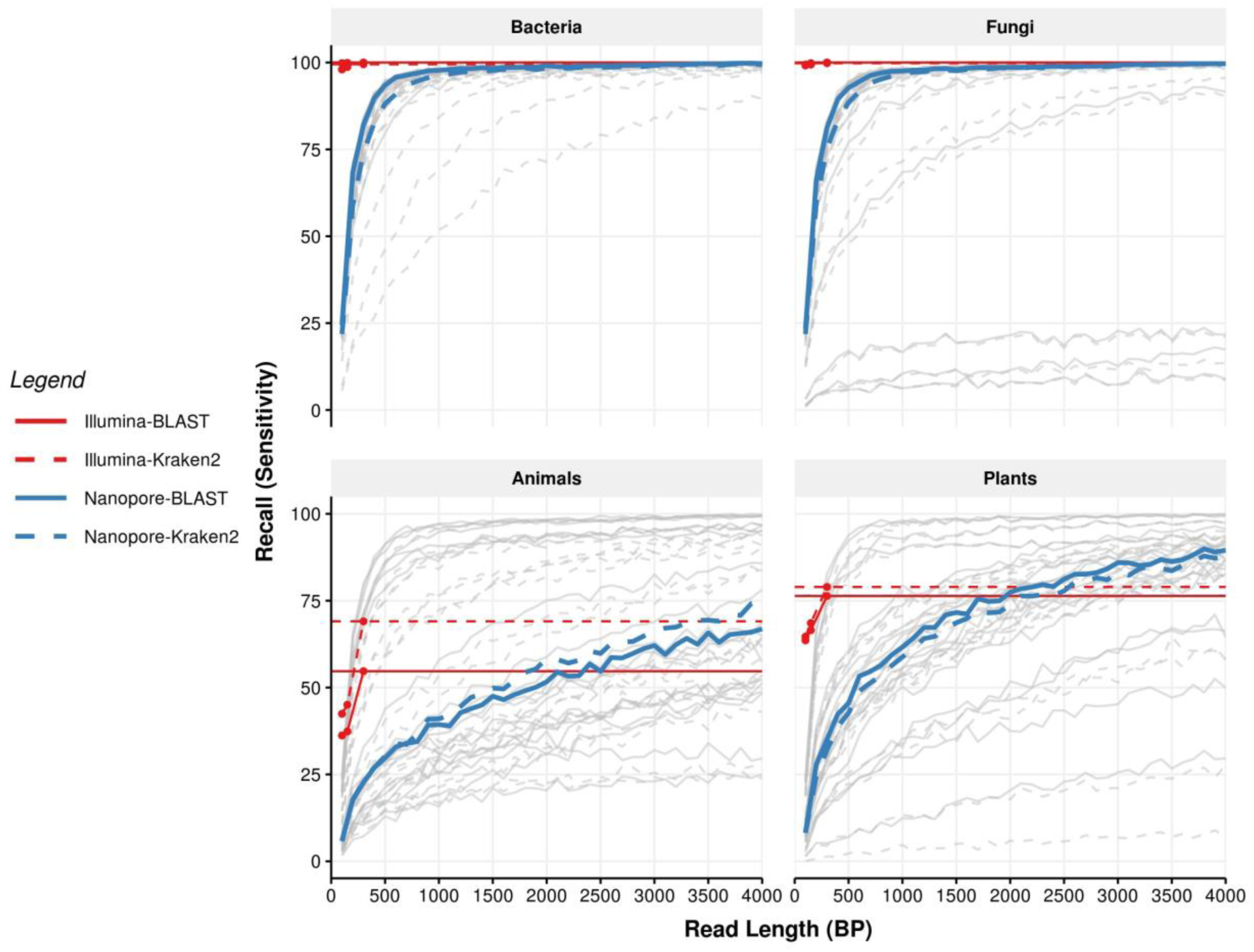
Recall at the family level. Each panel shows recall for the different kingdoms. Recall for Nanopore reads for individual taxa is indicated in grey, with median recall indicated by dashed lines, either blue (Kraken2) or red (BLAST). The recall rates for 300 bp Illumina reads are shown as thin solid lines, again either blue (Kraken2) or red (BLAST). Coloured points show the recall for all Illumina reads of all lengths (100 bp, 150 bp, and 300 bp).

**Supplementary Figure 3.**
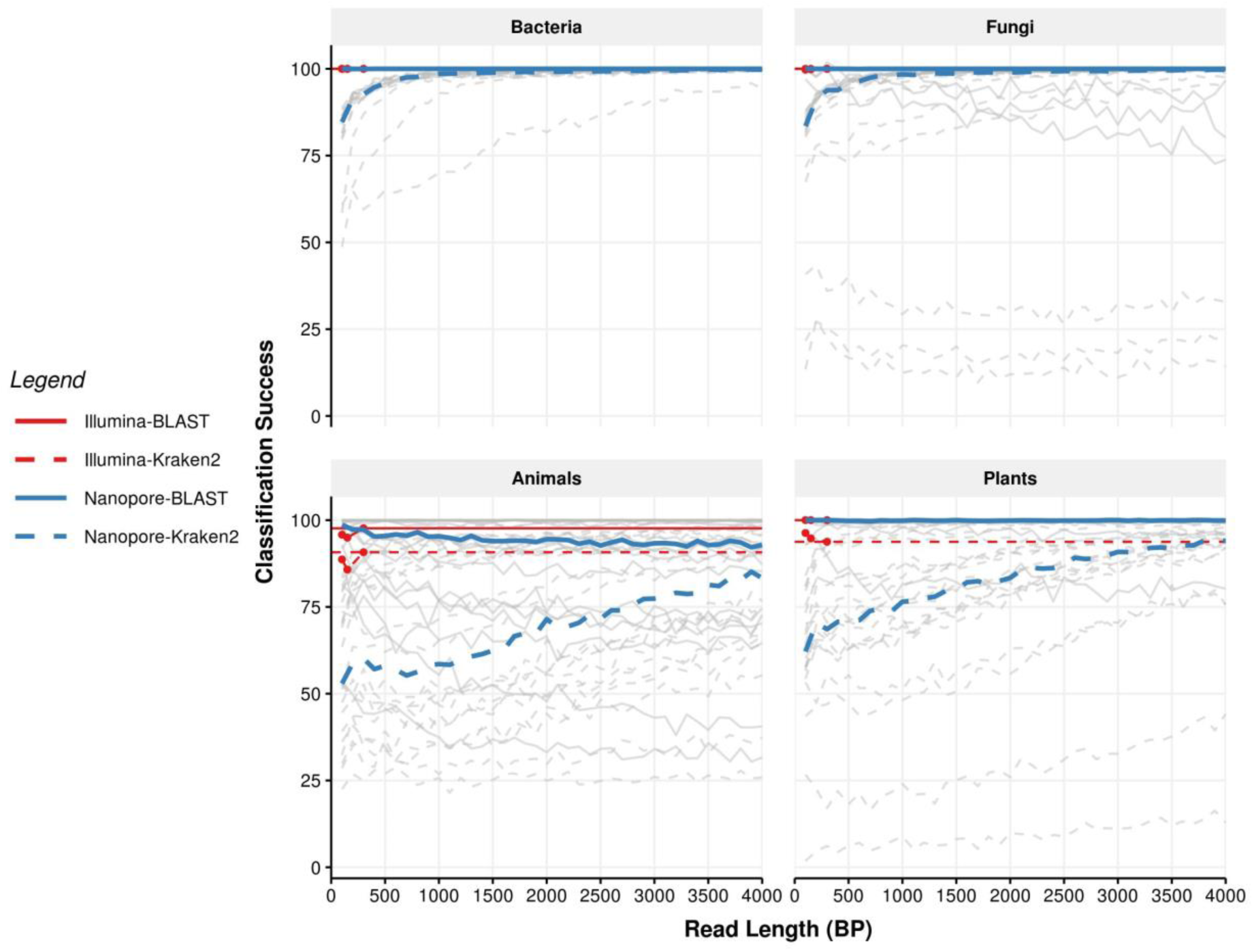
Classification success at the family level. Each panel shows classification success for the different kingdoms. Classification success for individual taxa is indicated in grey, with median classification success shown by solid lines (Illumina) or dashed lines (Nanopore). Blue indicates classification success rates for reads classified using Kraken2, while red indicates those classified using BLAST. For animal and plants, the classification success of Kraken2 depends strongly on read length, and never surpasses BLAST or Illumina at any length.

**Supplementary Table 1.** List of species include in the in silico mock community, with associated Kingdom and NCBI.

| Species | Kingdom | NCBI Accession |
| --- | --- | --- |
| *Actinidia chinensis* | Plantae | CM009654.1 |
| *Ananas comosus* | Plantae | CM003813.1 |
| *Arabidopsis thaliana* | Plantae | CP002684.1 |
| *Brassica nigra* | Plantae | CM004491.1 |
| *Camelina sativa* | Plantae | CM002729.1 |
| *Citrus sinensis* | Plantae | CM001701.1 |
| *Dioscorea rotundata* | Plantae | BDMI01000001.1 |
| *Eutrema salsugineum* | Plantae | CM001778.1 |
| *Gossypoides kirkii* | Plantae | CM008980.1 |
| *Leersia perrieri* | Plantae | CM002476.1 |
| *Malus domestica* | Plantae | CM007867.1 |
| *Micromonas* sp. | Plantae | CP001574.1 |
| *Panicum hallii* | Plantae | CM008046.2 |
| *Raphanus sativus* | Plantae | CM007999.1 |
| *Rosa chinensis* | Plantae | CM009582.1 |
| *Setaria italica* | Plantae | CM004364.1 |
| *Solanum lycopersicum* | Plantae | CM001064.3 |
| *Sorghum bicolor* | Plantae | CM000760.3 |
| *Theobroma cacao* | Plantae | LT594788.1 |
| *Trifolium pratense* | Plantae | LT555306.1 |
| *Amphiprion percula* | Animalia | CM009708.1 |
| *Bos indicus* | Animalia | CM003021.1 |
| *Capra hircus* | Animalia | CM001710.2 |
| *Chrysemys picta* | Animalia | CM002655.1 |
| *Columba livia* | Animalia | CM007525.1 |
| *Cyprinus carpio* | Animalia | LN590701.1 |
| *Drosophila busckii* | Animalia | CP012523.1 |
| *Equus caballus* | Animalia | CM000377.2 |
| *Falco peregrinus* | Animalia | CM007505.1 |
| *Homo sapiens* | Animalia | CM004593.1 |
| *Lycaon pictus* | Animalia | CM007565.1 |
| *Macaca mulatta* | Animalia | CM000308.1 |
| *Microcebus murinus* | Animalia | CM007661.1 |
| *Mus musculus* | Animalia | CM004154.1 |
| *Oncorhynchus tshawytscha* | Animalia | CM009202.1 |
| *Oryctolagus cuniculus* | Animalia | CM000790.1 |
| *Ovis aries* | Animalia | CM008472.1 |
| *Takifugu rubripes* | Animalia | HE602535.1 |
| *Timema cristinae* | Animalia | CM007794.2 |
| *Xiphophorus maculatus* | Animalia | CM008938.1 |
| *Agaricus bisporus* | Fungi | CP015470.1 |
| *Alternaria solani* | Fungi | CP022024.1 |
| *Colletotrichum higginsianum* | Fungi | CM004455.1 |
| *Cryptococcus gattii* | Fungi | CP025759.1 |
| *Debaryomyces hansenii* | Fungi | CR382133.2 |
| *Eremothecium sinicaudum* | Fungi | CP014242.1 |
| *Flammulina velutipes* | Fungi | CM002695.1 |
| *Fusarium verticillioides* | Fungi | CM000578.1 |
| *Kluyveromyces lactis* | Fungi | CR382121.1 |
| *Komagataella phaffii* | Fungi | LT962476.1 |
| *Lachancea nothofagi* | Fungi | LT598449.1 |
| *Malassezia sympodialis* | Fungi | LT671813.1 |
| *Milleriozyma farinosa* | Fungi | FO082059.1 |
| *Ogataea parapolyomorpha* | Fungi | CM002300.1 |
| *Saccharomyces cerevisiae* | Fungi | BK006935.2 |
| *Sporisorium scitamineum* | Fungi | CP010913.1 |
| *Trichoderma reesei* | Fungi | CP016232.1 |
| *Valsa mali* | Fungi | CM003098.1 |
| *Yarrowia lipolytica* | Fungi | HG934059.1 |
| *Zygosaccharomyces rouxii* | Fungi | CU928173.1 |
| *Acidithiobacillus ferrivorans* | Bacteria | LT841305.1 |
| *Bacillus thuringiensis* | Bacteria | CP015250.1 |
| *Bacillus velezensis* | Bacteria | CP025939.1 |
| *Bifidobacterium longum* | Bacteria | CP013673.1 |
| *Bordetella bronchiseptica* | Bacteria | CM002881.1 |
| *Brucella melitensis* | Bacteria | CP018494.1 |
| *Campylobacter jejuni* | Bacteria | CP012689.1 |
| *Caulobacter crescentus* | Bacteria | AE005673.1 |
| *Cellvibrio japonicus* | Bacteria | CP000934.1 |
| *Escherichia albertii* | Bacteria | AP014855.1 |
| *Gordonibacter* sp. | Bacteria | LT827128.1 |
| *Klebsiella pneumoniae* | Bacteria | CP025088.1 |
| *Mycobacterium tuberculosis* | Bacteria | CP023640.1 |
| *Ornithobacterium rhinotracheale* | Bacteria | CP006828.1 |
| *Pseudomonas arsenicoxydans* | Bacteria | LT629705.1 |
| *Salmonella enterica* | Bacteria | CP007400.2 |
| *Serratia symbiotica* | Bacteria | LN890288.1 |
| *Staphylococcus aureus* | Bacteria | CP012974.1 |
| *Treponema pallidum* | Bacteria | CP020366.1 |
| *Vibrio cholerae* | Bacteria | LT907989.1 |

